# Biophysical Basis of Hb-S Polymerization in Red Blood Cell Sickling

**DOI:** 10.1101/676957

**Authors:** W. Li

## Abstract

Sickle cell disease (SCD) is an autosomal recessive genetic disease caused by the Glu6Val mutation in the β chain (Hb) of the oxygen-carrying hemoglobin protein in sicklemia patients. In the molecular pathogenesis of SCD, the sickle hemoglobin (Hb-S) polymerization is a major driver for structural deformation of red blood cells, i.e. red blood cell (RBC) sickling. Biophysically, it still remains elusive how this SCD-linked E6V mutation leads to Hb-S polymerization in RBC sickling. Therefore, with a comprehensive set of analysis of experimental Hb structures, this letter highlights electrostatic repulsion as a key biophysical mechanism of Hb-S polymerization in RBC sickling, which provides atomic-level insights into the functional impact of the SCD-linked E6V substitution from a biophysical point of view.

**SIGNIFICANCE:** During the past 25 years, a total of 104 Hb-related structures have been deposited in PDB. For the first time, this article presents a comprehensive set of electrostatic analysis of the 104 experimental structures, highlighting electrostatic repulsion as a fundamental biophysical mechanism for Hb-S polymerization in RBC sickling. The structural and electrostatic analysis here also provides biophysical insights into the functional impact of the SCD-linked E6V substitution.

## INTRODUCTION

Sickle cell disease (sicklemia, SCD) is a severe hereditary form of anemia that results from a single genetic mutation at the sixth codon of the β-*globin* gene (1–4). The homozygous missense genetic mutation leads to a Glu6Val (E6V) substitution in the amino acid sequence of β globin (Hb), resulting in the sickle hemoglobin (Hb-S, α_2_β^S^_2_), while the oxygen-carrying hemoglobin protein is normally a tetramer in the form of α_2_β_2_ (5–9).

Among human genetic diseases, SCD is a rather rare case where a single mutation exhibits a wide range of effects at various levels, including molecular and cellular, both positive and negative (1, 2, 10, 11). However, from a structural and biophysical point of view, it still remains elusive how this sicklemia-linked E6V mutation leads to Hb-S polymerization in RBC sickling. The central aim of this article is to address this question with a comprehensive set of structural electrostatic analysis of all Hb-related structures in the Protein Data Bank (PDB) (12).

## MATERIALS AND METHODS

### Experimentally determined β-globin structures in PDB

To retrieve all experimentally determined β-globin structures as of June 18, 2019, a structure search was conducted into PDB (12) using three parameters: 1), Text Search = β-globin; 2), TAXONOMY = Homo sapiens; 3), Molecule = hemoglobin subunit β. As of June 18, 2019, there are 104 Hb-related structures in PDB. Among them, 97 were determined by X-ray crystallography, 2 by neutron diffraction (PDB IDs: 2DXM and 3KMF), 2 by solution NMR spectroscopy (PDB IDs: 2M6Z and 2H35), and 3 by Cryo-electron microscopy (Cryo-EM, PDB IDs: 5NI1, 6NBD and 6NBC). In total, the depositions of the 104 structures cover a time period of ~25 years, starting from October, 1993, ending in June, 2019. The 104 PDB IDs are 1A00, 1A01, 1A0U, 1A0Z, 1DXT, 1DXU, 1DXV, 1HDB, 1RQA, 1Y0T, 1Y0W, 2D5Z, 2D60, 2DN1, 2DN2, 2DN3, 2DXM, 2H35, 2HHB, 2HHE, 2M6Z, 2W6V, 2W72, 2YRS, 3B75, 3D17, 3D7O, 3DUT, 3HHB, 3HXN, 3IC0, 3IC2, 3KMF, 3NL7, 3NMM, 3ODQ, 3ONZ, 3OO4, 3OO5, 3P5Q, 3QJB, 3QJC, 3QJD, 3QJE, 3R5I, 3S65, 3S66, 3SZK, 3W4U, 3WCP, 3WHM, 4FC3, 4HHB, 4IJ2, 4L7Y, 4M4A, 4M4B, 4MQC, 4MQG, 4MQH, 4MQI, 4N7N, 4N7O, 4N7P, 4N8T, 4NI0, 4NI1, 4ROL, 4ROM, 4WJG, 4X0L, 4XS0, 5E29, 5E6E, 5E83, 5EE4, 5HU6, 5HY8, 5JDO, 5KDQ, 5KSI, 5KSJ, 5NI1, 5SW7, 5U3I, 5UCU, 5UFJ, 5URC, 5VMM, 5WOG, 5WOH, 5X2R, 5X2S, 5X2T, 5X2U, 6BB5, 6BNR, 6BWP, 6BWU, 6DI4, 6FQF, 6HAL, 6NBC, 6NBD.

### Structural analysis of the β-globin structures

While close-range hydrophobic interactions play an important part in biomolecular structure and function (13), intramolecular or intermolecular electrostatic interactions, such as salt bridge, hydrogen bond, and charge-dipole interaction, stabilize or de-stabilize biomolecules, too (14, 15). Here, the primary impact of the sicklemia-linked E6V mutation is an electrostatic one, i.e., the removal of Glu6’s negatively charged side chain, while a valine (with hydrophobic side chain) is installed at amino acid residue position 6 of Hb (1, 3, 4). After the retrieval of the 104 Hb structures from PDB, a set of structural electrostatic analysis was carried out as described in (16), including analysis of salt bridging (side chain alone) and hydrogen bonding (both main chain and side chain) networks. In particular, the two NMR ensembles (PDB IDs: 2M6Z and 2H35) were splitted into 40 standalone structures (16), since each NMR ensemble consists of 20 structural models. As a result, a total of 142 β-globin structural models were retrieved and extracted from PDB, as of June 18, 2019.

## RESULTS

### Structural and electrostatic analysis of Glu6 of Hb

For the 142 Hb structural models, a total of 12176 salt bridges were structurally identified with a cut-off distance at 4 Å(16). Specifically, only 4 side chain salt bridges (Tables 1 and 2 of Supporting Material **supps.pdf**) were structurally identified for Glu6 of Hb in the 142 models, which is considered statistically insignificant due to a ratio of only 2.82% (4/142) here. Furthermore, while Glu6 of Hb does form hydrogen bonds with other residues, no side chain hydrogen bond was identified for it (Tables 3-7 of Supporting Material **supps.pdf**). Taken together, the structural electrostatic analysis here does not align with the experimental observation that the SCD-linked E6V mutation, whose primary impact is the removal of Glu6’s negatively charged side chain, is inextricably linked to the molecular pathogenesis of SCD (1–4). This misalignment suggests that the net impact of the E6V substitution is probably an indirect one, i.e., the loss of Glu6’s side chain leads to indirect local electrostatic equilibrium perturbation, which is of functional significance for the overall structural stability of Hb, because Glu6 itself does not seem to be engaged in any significant electrostatic interaction with Hb’s interacting partners, according to the experimental structural investigations during the past 25 years.

**Table 1:** The salt bridges formed between Glu7 and Lys132 in the second structural model (PDB ID: 2M6Z).

| PDB | Residue A | Atom A | Residue B | Atom B | Inter-atomic distance (Å) |
| --- | --- | --- | --- | --- | --- |
| 2M6Z-2 | B_LYS_132 | NZ | B_GLU_7 | OE1 | 2.898 |
| 2M6Z-2 | B_LYS_132 | NZ | B_GLU_7 | OE2 | 3.967 |
| 2M6Z-2 | D_LYS_132 | NZ | D_GLU_7 | OE1 | 2.891 |
| 2M6Z-2 | D_LYS_132 | NZ | D_GLU_7 | OE2 | 3.959 |

**Table 2:**
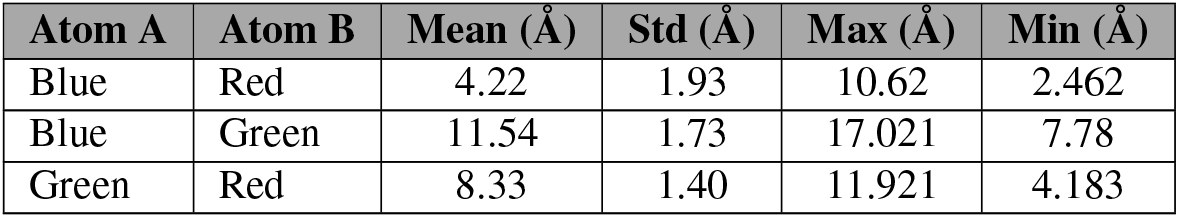
The spatial distances between the three types of atoms of the Glu6-Glu7-Lys132 axis (Fig 2). In this table, the details of the three colors (Blue, Red and Green) are provided in the caption of Fig 2

### Structural and electrostatic analysis of Glu7 of Hb

To address this misalignment, the amino acid sequence and the three-dimensional structure of Hb (with or without its partners) were analyzed with a careful inspection, in addition to an automated residue pair-specific ranking of the numbers of all 12176 salt bridges, after which Glu7 of Hb stood out because,

1. Both sequence-wise and structurally, Glu7 of Hb is the closest neighbour of Glu6 of Hb.
2. Glu7 and Glu6 are of the same residue type, both carry side chains with negative electric charge.
3. In the residue pair-specific ranking, one particular residue pair stood out, i.e., Glu7 and Lys132 of Hb, where 936 side chain salt bridges were identified for the 142 Hb structural models, ranking it the second largest, according to Table 2 of Supporting Material **supps.pdf**.
4. Glu7 did not form any other salt bridge, according to Table 2 of Supporting Material **supps.pdf**.

In addition to the 936 Lys132-Glu7 salt bridges, Lys132 only forms 7 side chain salt bridges with Asp136, which is considered statistically insignificant due to a ratio of only 4.92% (7/142), too. To this end, the salt bridge analysis here established an axis of the amino acid residue triplet, i.e., Glu6-Glu7-Lys132, for which the result of the hydrogen bonding analysis is statistically insignificant and consequently irrelevant here, according to Table 7 in Supporting Material **supps.pdf**.

### Structural electrostatic insights into the Glu6-Glu7-Lys132 axis

Structural stability-wise, of particular interest is the Glu7-Lys132 salt bridges (Table 1, Fig 1), which constitutes an electrostatic clip (16) that holds the two α helix fragments (cyan and green, Figs 1) together, and plays a positive role in the overall structural stability of Hb, either apo or bound to its interacting partners. From Table 1 and Fig 1, it can be seen clearly that Glu7 and Lys132 of Hb formed four intra-chain structurally stabilizing salt bridges (i.e., four strong electrostatic clips (16)), including chains B and D (PDB ID: 2M6Z), to keep the tetramer (α_2_β_2_) (5–9) hemoglobin from falling apart in human blood.

**Figure 1:**
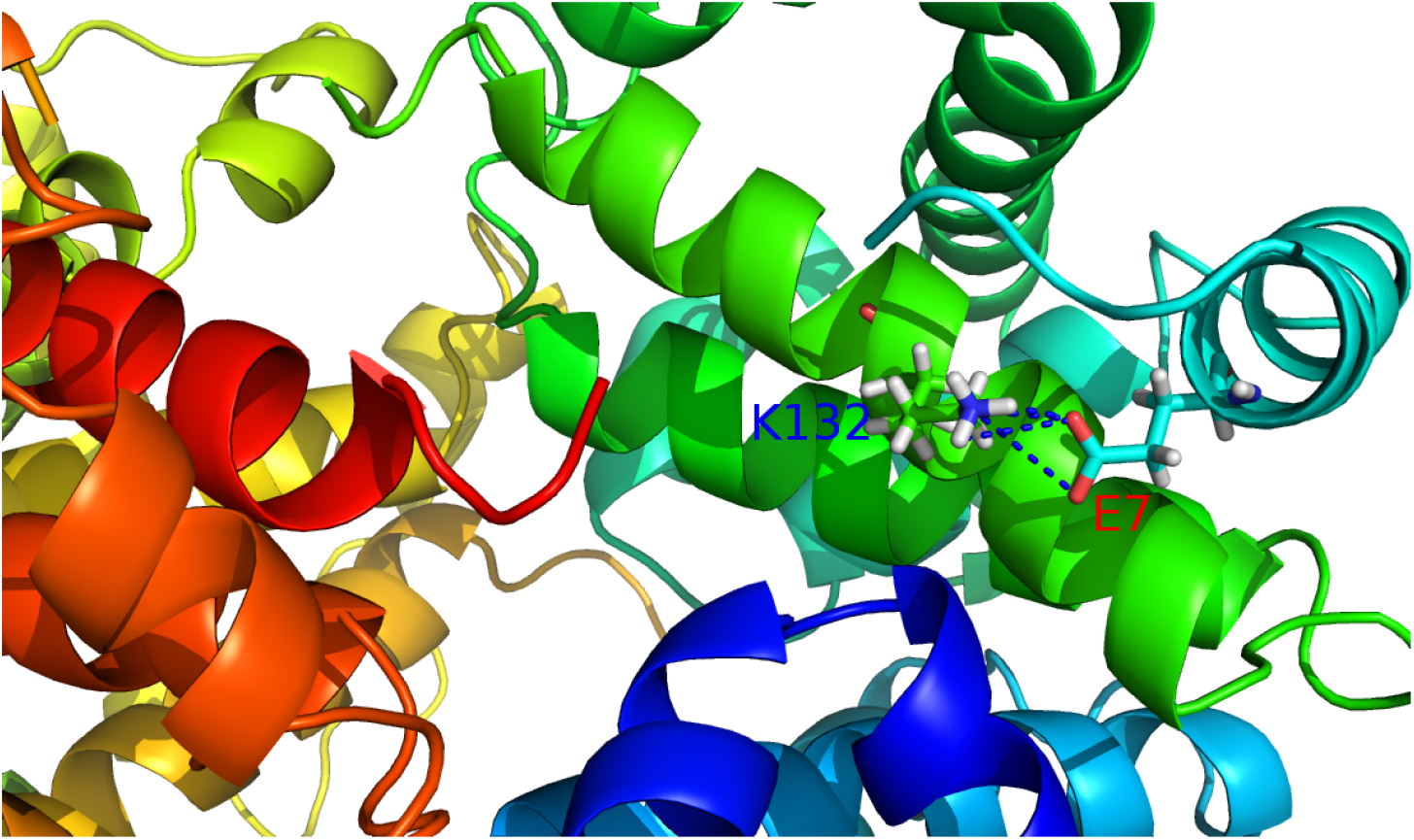
The Glu7-Lys132 salt bridge in the second model (PDB ID: 2M6Z, Table 1).

### Geometric and biophysical insights into the Glu6-Glu7-Lys132 axis

To characterize the structural and functional consequence(s) of the SCD-linked E6V substitution, the three-dimensional geometric configurations of the Glu6-Glu7-Lys132 axis are projected into a two-dimensional plane according to the experimentally measured atomic coordinates, as shown below in Fig 2 and Table 2.

**Figure 2:**
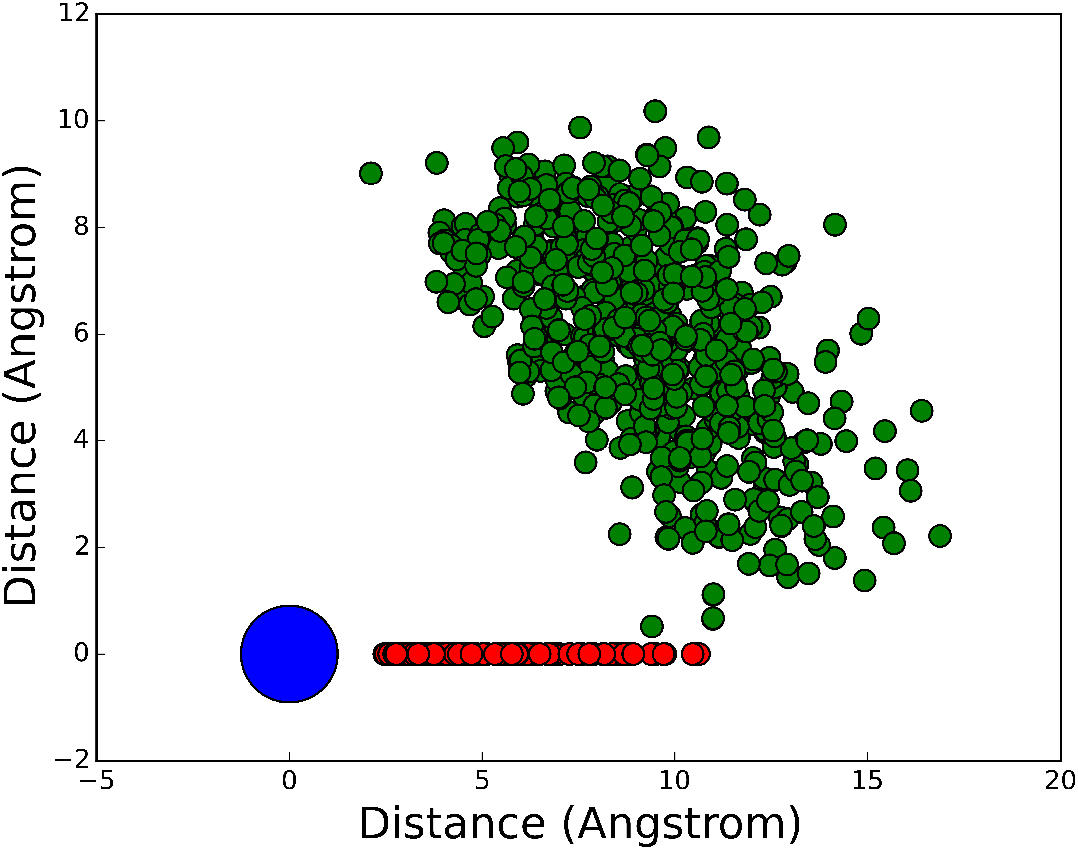
Two-dimensional projection of the three-dimensional geometric configurations of the Glu6-Glu7-Lys132 axis. In this figure, the large black solid circle represents the N_ζ_ atom of Lys132, the small red and green solid circles represent the side chain oxygen atoms (O_ε1_ and O_ε2_) of Glu7 and Glu6, respectively. The inter-actomic distances are included in Table 2.

With a visual inspection of Fig 2, it is quite clear that the negatively charged side chain of Glu6 forms an electric shield (the green solid circles in Fig 2) for the intra-chain side chain salt bridge between Glu7 and Lys132, which plays a crucial role in the overall structural stability of the tetramer (α_2_β_2_) (5–9) hemoglobin.

1. In case any positively charged particle (atoms for instance) in the blood approaches the Lys132-Glu7 salt bridge, the existence of negatively charged side chain of Glu6 (the green solid circles in Fig 2) constitute an electrostatic attractive force for the positively charged particle, as it is energetically favourable for the positively charged particle to approach Glu6, instead of the salt-bridged Glu7, because the negatively charged side chain of Glu6 is mostly exposed to the solvent, according to Tables 1 and 2 of Supporting Material **supps.pdf**. Thus, Glu6’s negatively charged side chains act as a electric shield (via electrostatic attraction) to towards the structural stability of the α_2_β_2_-stabilizing Lys132-Glu7 salt bridge, because a positively charged particle will decrease the strength of the Lys132-Glu7 salt bridge via competitive electrostatic interaction with Glu7’s side chain, if it does not break the α_2_β_2_-stabilizing Lys132-Glu7 salt bridge.
2. In case any negatively charged particle in the blood approaches the Lys132-Glu7 salt bridge, the existence of negatively charged side chain of Glu6 (the green solid circles in Fig 2) constitute an electrostatic repulsive force against the approaching negatively charged particle. Thus, Glu6’s negatively charged side chains act as a electric shield (via electrostatic repulsion) towards the structural stability of the α_2_β_2_-stabilizing Lys132-Glu7 salt bridge.

### WHY IS THE ELECTROSTATIC REPULSION MECHANISM SO IMPORTANT HERE?

In addition to the E6V mutant β-globin (Hb-S), a different form of sickle hemoglobin (Hb-S, Lys132Asn, K132N) was also reported in 2004 in (17), where the Lys132-Glu7 salt bridge is disrupted as Asn132 does not carry a charged side chain, and Asn132 is replaced by Lys132, which carries a positively charged side chain. Structurally, the net effect of this Asn-Lys substitution is the removal of the α_2_β_2_-stabilizing Lys132-Glu7 salt bridge, highlighting the perturbation of local electrostatic equilibrium by SCD-linked mutations such as K132N and E6V, too.

Previous molecular dynamics simulation studies also suggest that Hb-S polymerization is favoured by inter-facial electrostatic interactions (18), and is inhibited when subjected to hypoxic (low pH environment) conditions, where the side chain of Glu6 gets neutralized and the electrostatic repulsion mechanism disappears. This molecular dynamics simulation study provides a direct support that the electrostatic repulsion mechanism, previously described in (16, 19, 20), is a key biophysical basis for Hb-S polymerization in RBC sickling, too.

To further test this electrostatic repulsion mechanism, the 142 Hb structural models were imported into the PROPKA software package (21). Afterwards, the protonation states/proton occupancies (*θ*) of all Glu6 side chains were calculated with the pKa values (3.897 ± 0.645) as an input for the rearranged classic Henderson-Hasselbalch equation (22). Given the physiological pH (7.4) of human blood, only 0.009 ± 0.062 % of the Glu6 residues remain protonated, i.e., neutral, and 0.991 ± 0.062 of the Glu6 residues remain deprotonated, i.e., negatively charged. The protonation states of Glu6 residues allow them to form negatively charged electric shields to ensure the stability of the Glu7-Lys132 salt bridge against the approach of negatively charged particles in human blood. In particular, the two β chains carry two Glu6 residues (chains B and D) with negatively charged solvent-exposed side chains, too. If it is not for the existence of the Glu6 as an electric shield (Fig 2 and Table 2), it will be energetically more favorable for Hb-S to polymerize because the electrostatic repulsion mechanism disappears due to the SCD-linked E6V substitution, and thus making it easier for the two β chains fuse into larger biomolecular complexes/particles/chemical groups (20). Taken together, it is due to the protonation state of the Glu6 side chain at physiological pH that the SCD-linked E6V substitution abolishes the electrostatic repulsion mechanism as a valine does not carry a charged side chain, making it easier for Hb-S (E6V) molecules to approach each other and fuse (20) with each other and polymerize to cause RBC sickling(23, 24).

Moreover, the pKa values of all Glu7 side chains were also calculated with the PROPKA software package (21) to be 3.897 ± 0.645. Given a physiological pH (7.4) of human blood, 0.001 ± 0.002 % of the Glu7 residues remain protonated, i.e., neutral, and 0.999 ± 0.002 of the Glu7 residues remain deprotonated, i.e., negatively charged, such that its side chain is able to form a structurally stable and α_2_β_2_-stabilizing salt bridge with the side chain of Lys132 (Fig 1). Furthermore, it is also due to the existence of a rather stable Lys132-Glu7 salt bridge (Fig 1) that the pKa of Glu7 is clearly lower that its intrinsic value (4.2), while the pKa of Glu6 is closer to 4.2 due to the absence of a rather stable salt bridge or hydrogen bond, which supports the overall performance of the pKa calculation of the PROPKA software package (21) here.

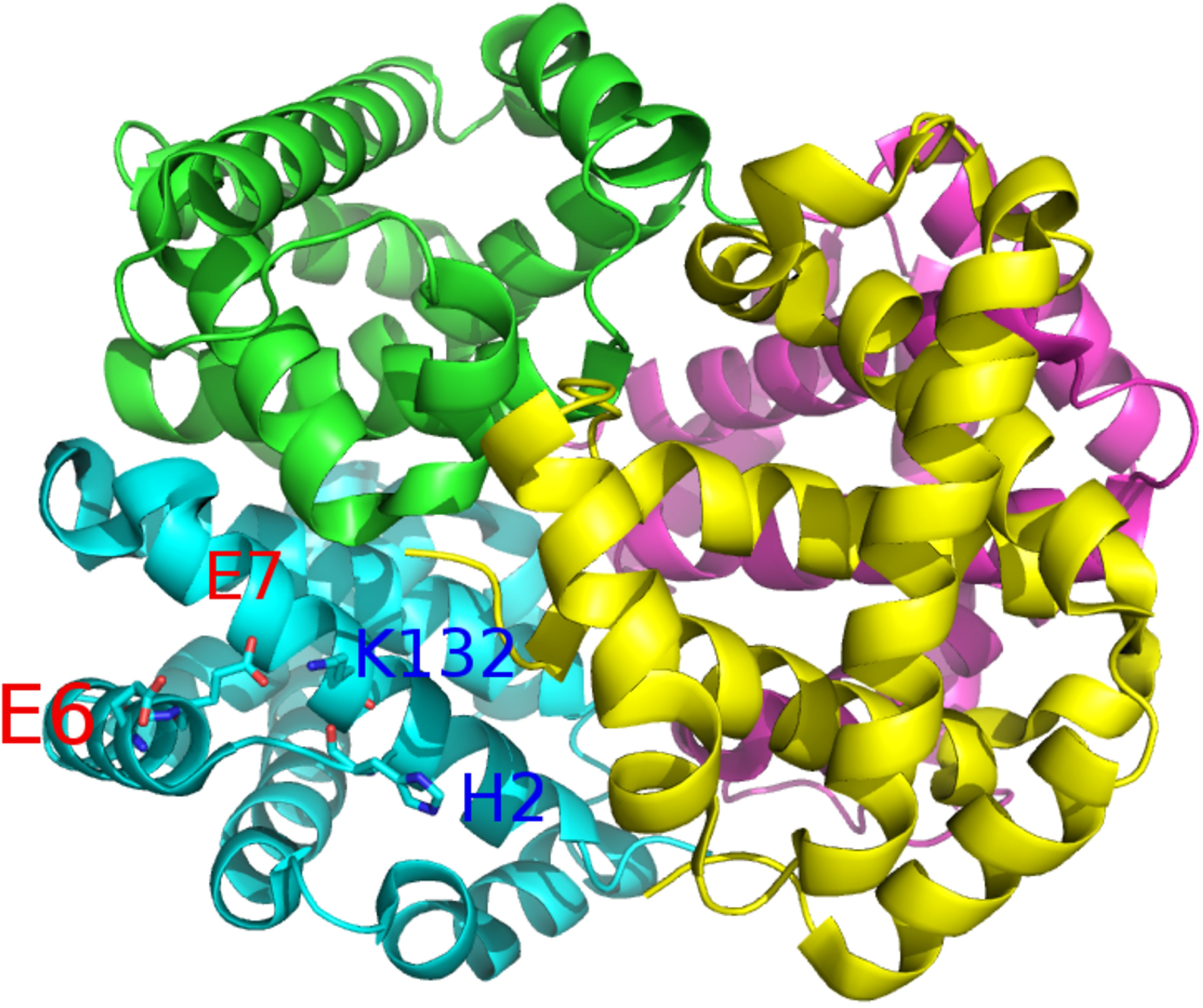

## REFERENCES

1. Pauling, L., H. A. Itano, S. J. Singer, and I. C. Wells, 1949. Sickle Cell Anemia, a Molecular Disease. Science 110:543–548.

2. Kato, G. J., F. B. Piel, C. D. Reid, M. H. Gaston, K. Ohene-Frempong, L. Krishnamurti, W. R. Smith, J. A. Panepinto, D. J. Weatherall, F. F. Costa, and E. P. Vichinsky, 2018. Sickle cell disease. Nature Reviews Disease Primers 4:18010.

3. Johannesen, E., and V. Nguyen, 2018. Massive Ovarian Edema in a Girl with Hemoglobin SC Disease. Case Reports in Pathology 2018:1–3.

4. Demirci, S., N. Uchida, and J. F. Tisdale, 2018. Gene therapy for sickle cell disease: An update. Cytotherapy 20:899–910.

5. Parrow, N. L., H. Tu, J. Nichols, P.-C. Violet, C. A. Pittman, C. Fitzhugh, R. E. Fleming, N. Mohandas, J. F. Tisdale, and M. Levine, 2017. Measurements of red cell deformability and hydration reflect HbF and HbA 2 in blood from patients with sickle cell anemia. Blood Cells, Molecules, and Diseases 65:41–50.

6. Oksenberg, D., K. Dufu, M. P. Patel, C. Chuang, Z. Li, Q. Xu, A. Silva-Garcia, C. Zhou, A. Hutchaleelaha, L. Patskovska, Y. Patskovsky, S. C. Almo, U. Sinha, B. W. Metcalf, and D. R. Archer, 2016. GBT440 increases haemoglobin oxygen affinity, reduces sickling and prolongs RBC half-life in a murine model of sickle cell disease. British Journal of Haematology 175:141–153.

7. Kim, H. C., 2014. Red cell exchange: special focus on sickle cell disease. Hematology 2014:450–456.

8. Cohen, S. I. A., and L. Mahadevan, 2013. Hydrodynamics of Hemostasis in Sickle-Cell Disease. Physical Review Letters 110.

9. Dufu, K., M. Patel, D. Oksenberg, and P. Cabrales, 2018. GBT440 improves red blood cell deformability and reduces viscosity of sickle cell blood under deoxygenated conditions. Clinical Hemorheology and Microcirculation 70:95–105.

10. Aidoo, M., D. J. Terlouw, M. S. Kolczak, P. D. McElroy, F. O. ter Kuile, S. Kariuki, B. L. Nahlen, A. A. Lal, and V. Udhayakumar, 2002. Protective effects of the sickle cell gene against malaria morbidity and mortality. The Lancet 359:1311–1312.

11. Aprelev, A., W. Stephenson, H. M. Noh, M. Meier, and F. A. Ferrone, 2012. The Physical Foundation of Vasoocclusion in Sickle Cell Disease. Biophysical Journal 103:L38–L40.

12. Berman, H., K. Henrick, and H. Nakamura, 2003. Announcing the worldwide Protein Data Bank. Nature Structural & Molecular Biology 10:980–980.

13. Chu, X., Y. Wang, L. Gan, Y. Bai, W. Han, E. Wang, and J. Wang, 2012. Importance of Electrostatic Interactions in the Association of Intrinsically Disordered Histone Chaperone Chz1 and Histone H2A.Z-H2B. PLoS Computational Biology 8:e1002608.

14. Nakamura, H., 1996. Roles of electrostatic interaction in proteins. Quarterly Reviews of Biophysics 29:1–90.

15. Kumar, S., and R. Nussinov, 2002. Close-Range Electrostatic Interactions in Proteins. ChemBioChem 3:604–617.

16. Li, W., 2017. How do SMA-linked mutations of *SMN1* lead to structural/functional deficiency of the SMA protein? PLOS ONE 12:e0178519.

17. Luo, H., A. H. Adewoye, S. H. Eung, T. P. Skelton, K. Quillen, L. McMahon, M. H. Steinberg, and D. H. K. Chui, 2004. A Novel Sickle Hemoglobin: Hemoglobin S-South End. Journal of Pediatric Hematology/Oncology 26:773–776.

18. Papageorgiou, D. P., S. Z. Abidi, H.-Y. Chang, X. Li, G. J. Kato, G. E. Karniadakis, S. Suresh, and M. Dao, 2018. Simultaneous polymerization and adhesion under hypoxia in sickle cell disease. Proceedings of the National Academy of Sciences 115:9473–9478.

19. Li, W., and G. Shi, 2019. How Ca_V_1.2-bound verapamil blocks Ca^2+^ influx into cardiomyocyte: Atomic level views. Pharmacological Research 139:153–157.

20. Li, W., 2016. Characterising the interaction between caenopore-5 and model membranes by NMR spectroscopy and molecular dynamics simulations. PhD thesis, University of Auckland

21. Olsson, M. H. M., C. R. Søndergaard, M. Rostkowski, and J. H. Jensen, 2011. PROPKA3: Consistent Treatment of Internal and Surface Residues in Empirical pKa predictions. Journal of Chemical Theory and Computation 7:525–537.

22. Li, W., 2017. Gravity-driven pH adjustment for site-specific protein pKa measurement by solution-state NMR. Measurement Science and Technology 28:127002.

23. Sundd, P., M. T. Gladwin, and E. M. Novelli, 2018. Pathophysiology of Sickle Cell Disease. Annual Review of Pathology: Mechanisms of Disease 14.

24. Hosseinzadeh, V. A., C. Brugnara, and R. G. Holt, 2018. Shape oscillations of single blood drops: applications to human blood and sickle cell disease. Scientific Reports 8.

